# HyenaSET: Hyena Sound Event Transcripts and benchmark animal2vec performance for parsing animal communication

**DOI:** 10.64898/2026.06.14.732108

**Authors:** Jana M. Woerner, Céline Angonin, Andrew S. Gersick, Kay E. Holekamp, Frants H. Jensen, Mark P. Johnson, Marsden H. M. Onsare, Malit O. Pioon, Julian C. Schäfer-Zimmermann, Ari Strandburg-Peshkin, Eli D. Strauss

**Author notes:** **Corresponding author:** Jana Woerner.

## Abstract

Here, we present HyenaSET, a large (~1640 hours) bioacoustic dataset derived from collar-mounted audio recorders deployed on 19 spotted hyenas in the Maasai Mara National Reserve, Kenya. Within this dataset, 243 hours have been strongly labeled by identifying the onset and offset of all vocalizations as well as their types, and the labels have been validated by experts on hyena vocalizations and behavior. Within the strongly labeled data, the total amount of time hyenas were vocalizing was 9.5 hours (3.9%). Furthermore, each vocalization has been manually identified as “focal” (emitted by the hyena wearing the collar) or “non-focal” (emitted by a nearby conspecific), making use of information from collar-mounted accelerometers that picked up vibrations of the animal’s throat when it produced vocalizations. In addition to the labeled data, we also provide a large corpus of unlabeled data from the same recordings, which can be used for un- or self-supervised machine learning tasks. To ensure reproducibility of this dataset as a benchmark in machine learning studies, we present it alongside five stratified cross-validation train/test splits to enable accurate comparisons, and we also provide a train/test split in which specific individuals are left out of the training set to assess generalizability across individuals. Finally, as a performance benchmark, we present baseline results for this dataset using *animal2vec*, a recently developed transformer-based model optimized for bioacoustic data.

## Background & Summary

Bioacoustics is a challenging but important component of multiple biological disciplines, relevant broadly across taxa.^1^ Animal communication plays a critical role in fundamental aspects of animal behavior such as courtship, cognition, parental care, social behavior, territorial defense, and cooperation.^2^ Bioacoustic data are typically collected by recording an observed animal with handheld directional microphones, passive acoustic monitors mounted in an environment, or animal-mounted recording tags.^3–5^ Once armed with recordings of animal sounds, playback studies can be used to interrogate call function across these contexts and test links between call structure and function.^4^ Biodiversity studies can leverage acoustic species classification to measure species occurrence.^5^ Research into the evolution and mechanisms underlying human spoken language often examines vocal production learning in animals.^6^ Conservation work on anthropogenic acoustic effects examines the ways that anthropogenic environments or sound sources may interfere with or otherwise modify animal communication.^7^ In sum, acoustic monitoring has wide applications across biology, and these applications require collecting and extracting information from bioacoustic data.

Drawing biological meaning from bioacoustic datasets typically requires identifying signals of interest from raw acoustic data streams. Bioacoustics researchers have often labeled their data manually by listening through audio recordings and/or visually inspecting spectrograms. As in many other scientific and societal realms, advances in machine learning are currently revolutionizing our ability to process bioacoustic datasets automatically or semi-automatically, opening up exciting new opportunities to analyze large-scale acoustic datasets that would otherwise have been infeasible to process.^8,9^ Yet, compared to datasets in other domains such as images,^10^ human speech,^11^ and text, ^12^ bioacoustic datasets are typically highly unbalanced, requiring a tremendous amount of person-time to curate datasets and generate ground truth information.^8^ Moreover, many bioacoustic datasets offer ground truth information only on the presence or absence of a given species or call in a given audio clip (this is called *weakly labeled*), rather than detailed information on the onsets and offsets of all acoustic events (*strongly labeled*). The high sparsity common within bioacoustic datasets means that it is challenging to find appropriate datasets for developing and testing modern machine learning approaches that require larger amounts of (labeled and unlabeled) data and temporally defined events – a constraint that limits the adaptation of modern machine learning methods to the bioacoustic domain.^8^

Here, we present a large (~1640 hours) bioacoustic dataset derived from collar-mounted audio recorders deployed on 19 spotted hyenas *(Crocuta crocuta)* in the Maasai Mara National Reserve, Kenya. By combining extensive continuous recordings with expert-validated, strongly labeled vocalizations, this dataset provides an extensive ground-truth dataset to train and validate machine learning models for automatically parsing animal communication signals, cross-species foundation models, and other methods that in turn can accelerate high-throughput animal behavior and communication studies. We present this dataset in a format that is designed to support the development and evaluation of modern machine learning approaches.

## Methods

### Study Species

Spotted hyenas live in large social groups (“clans”) and produce a wide variety of calls in various social contexts (Table 1). The clan is structured by strict dominance hierarchies, where social rank determines access to resources, and position in the social hierarchy is maintained through dyadic and coalitionary agonistic interactions that are often accompanied by vocal calls.^13–15^ Each clan defends a shared territory and contains multiple unrelated matrilines of females and their offspring, as well as one or more immigrant males.^13,14^ Female hyenas are philopatric, while most male hyenas disperse from their natal clan to seek mates in a new clan around ages 2-6.^16,17^ Hyena societies have high fission-fusion dynamics: though clans may encompass up to 130 members, individuals spend most of their time alone or in small subgroups that change composition throughout the day.^14,18,19^ Long-distance whoop calls allow hyenas to signal their location to distant group-mates, and sometimes, to rapidly gather to engage in cooperative behaviors such as border patrols or cooperative mobbing^20,21^. These fluid spatial dynamics allow group mates to avoid feeding competition with each other while also benefiting from communal resource defense against neighboring hyena clans and other competitors (i.e., lions).^14,19^ Spotted hyenas are efficient scavengers and hunters with a very flexible diet, allowing them to exploit both carrion and live prey, feeding on whichever prey species is most abundant or easiest to catch at any given time.^22,23^ Reproductive hyena mothers keep their young at a communal den, which functions as the social hub of the clan: young hyenas meet other group members that visit the den, practice the locomotor skills necessary for hunting and mating through play behavior, and learn their social rank within the larger group through repeated aggressive and submissive behaviors.^14,24,25^ Because of the social role of the den, this is a site of frequent vocal activity.

**Table 1.** Vocalization types labeled within the dataset.

| Call Type | Characteristics | Biological Context |
| --- | --- | --- |
| Whoop<sup>28</sup> | Low to mid frequency, tonal<br>Occur in bouts of two or more whoops<br>Initial and terminal whoop bout structure differs from those in the rest of the bout<br>Contain symmetrical or asymmetrical frequency sweeps | Long-distance calls used to quickly spread information across spatially dispersed social groups<sup>14</sup><br>May be used to recruit groupmates to a location or to advertise an individual's location/presence in the clan<sup>29</sup><br>Encode important information including the caller's age, sex, and identity<sup>30,31</sup> |
| Giggle | High-frequency, rapid frequency modulation | Emitted when a hyena is excited or fearful, such as when threatened, chased, or attacking others<sup>14,32</sup> |
| Squeal | Loud scream, often descending in frequency | Emitted during aggressive encounters when bitten or attacked |
| Growl | Low-frequency, noisy | Emitted during aggressive encounters when bitten or attacked |
| Groan | Low-frequency, tonal | "Come hither" vocalizations<br>Often used during affiliative behaviors, such as greeting ceremonies, and by mothers to coax their young out of den holes or to encourage them to follow her |
| Alarm rumble | Very low frequency, pulsed, extended | Emitted to warn others of possible dangers |
| Squitter | High frequency, extended<br>Only labeled as non-focal | Emitted by cubs to beg for food or milk from their mothers |
| Feeding | Broadband, noisy, rhythmic | Chewing and bone crunching noises emitted while eating |
| Snores | Low-frequency, rhythmic | Emitted while sleeping |
| Other |  | Includes small vocalizations that do not match other types, hybrid/fusion calls, and rare calls (lows) |

### Data Collection

This dataset was collected for the Communication & Coordination Across Scales (CCAS) Project, in conjunction with the Mara Hyena Project. CCAS is a multi-disciplinary project broadly focused on how communication drives coordination and decision-making in animal societies. The audio dataset presented here derives from 19 tracking collars deployed on members 20 months of age and older (age range: 23 months to 14.6 years) in a single clan of spotted hyenas residing in the Maasai Mara National Reserve, Kenya—the total number of individuals above this age threshold in the group was 23. These collars recorded audio continuously from January 1 until January 28, 2023, though most collars stopped recording between January 21-23. Due to memory limitations, a small number of collars (3 individuals) were programmed to record only during the night when hyenas are most active. From each collar, audio (32 kHz), accelerometer (1000 Hz), and magnetometer (50 Hz) data were recorded using a modified sound and movement recording DTAG.^3,26^ The DTAG was connected with a serial cable to a Gipsy 5 GPS logger (Technosmart Ltd, Rome, Italy) that recorded fine-scale location data. The PPS and timing information from the GPS logger were processed by the DTAG to enable time synchrony. The DTAG and Gipsy-5 were housed in a custom-designed housing with a GORE acoustic vent to protect the microphone from the environment. The recording unit was mounted on a Tellus Medium base collar with Iridium connectivity (Followit Sweden AB, Lindesberg, Sweden). To deploy collars, all hyenas were immobilized by a Kenya Wildlife Service-approved veterinarian (M. Onsare). At the end of the recording period, a hand-held radio transmitter was used to trigger a remote release system from the collar, though occasionally a few hyenas (n = 6) had to be sedated to recover the collar. This work followed ASM guidelines^27^ and was approved by the IACUC at Michigan State University (PROTO202200047) and Kenya Wildlife Service (KWS/904 to KEH).

### Data annotation and validation

Within this dataset, 243 hours have been strongly labeled by identifying the onset and offset of all vocalizations as well as the type of vocalization. Annotations were generated by trained human analysts, and all annotations were verified by J. Woerner and E. Strauss. Annotators used custom Matlab tools developed by M. Johnson and F. H. Jensen. This GUI integrated time-synchronized audio and accelerometer data (RMS energy after high-pass filtering with a 6-pole 50-Hz Butterworth filter) and visualized data in 10s periods for interactive labeling (Figure 1). Seven types of hyena vocalizations were annotated, including whoop, groan, alarm rumble, squeal, giggle, squitter, and growl (Table 1; Figure 2). Each annotation was designated as focal (emitted by the hyena wearing the collar) or non-focal (emitted by a nearby conspecific) based on the occurrence of throat vibrations detected on the high sample rate accelerometer. In cases where it was not possible to determine if a call was focal or non-focal, that missing information was labeled “unf” for “unknown focal.” Note that because squitters (nursing cries) are produced only by juveniles and only adults were wearing collars, all squitters were labeled as non-focal. In addition to the above hyena vocalizations, feeding noises were also annotated for the purposes of detecting feeding events. Snores were annotated to prevent confusion between groans and snores as they share some similar sound characteristics. We used an “other” call category to capture hyena sounds that did not fall clearly into any of the above categories, that seemed to be intermediates between two or more call types (hybrid) or consist of two or more call types joined together (fusion), or were calls that were extremely rare (e.g., lows). A number of other animal and non-animal sounds can be heard on these recordings, including zebras, hippos, crocodiles, lions, multiple bird and insect species, humans, and vehicles – none of these sounds were annotated.

**Figure 1.**
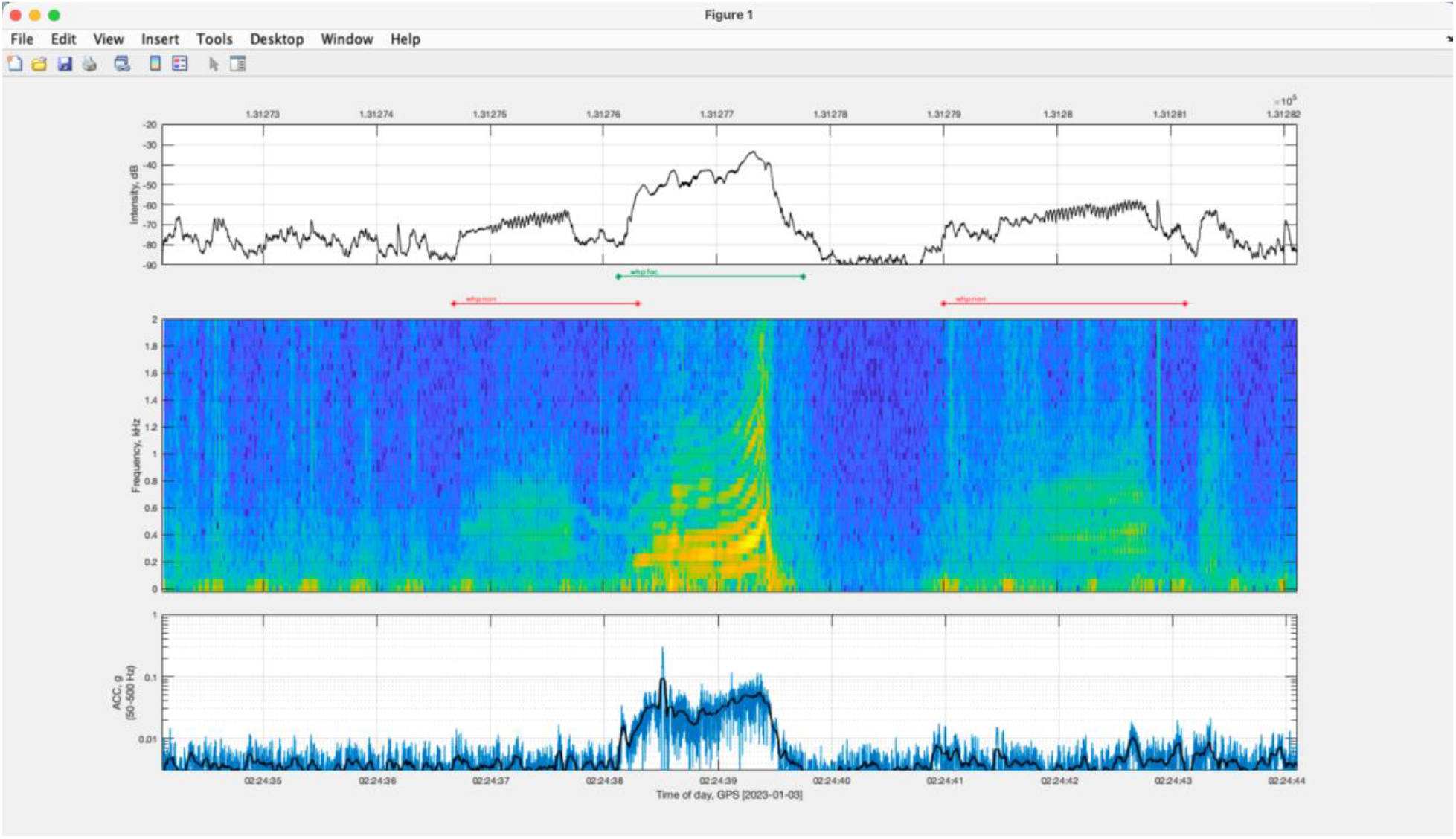
Matlab interface used by trained human analysts to generate annotations (developed by M. Johnson & F. H. Jensen). The top row shows sound intensity, the middle row shows the spectrogram, and the bottom row shows the intensity of vibrations detected by the accelerometer. Here, two non-focal whoops and one focal whoop are displayed.

**Figure 2.**
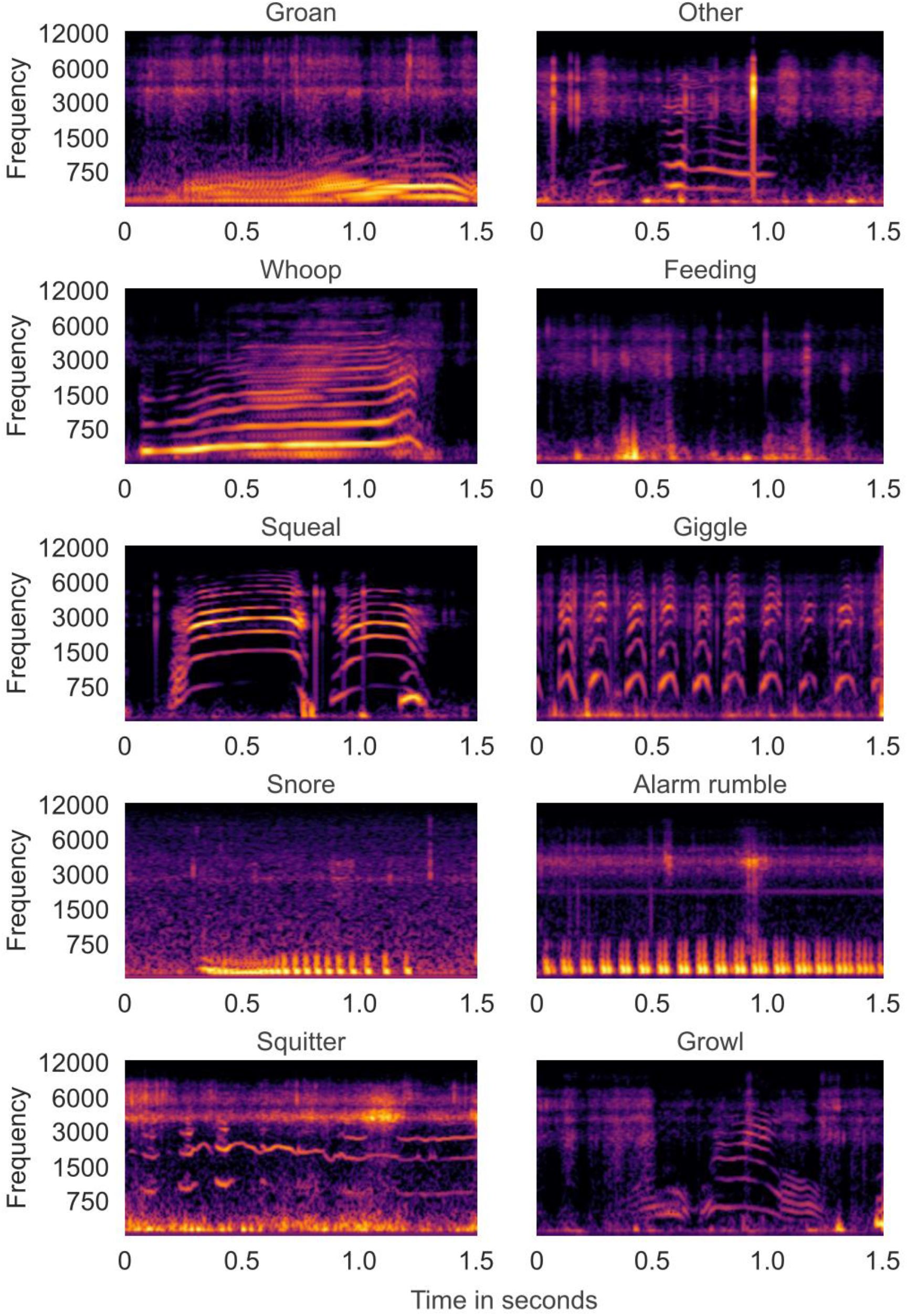
Mel spectrograms in dBr scale of clean representative examples for all classes in the HyenaSET dataset.

## Data Record

HyenaSET is released as 590,480 10-second samples, totaling approximately 1640 h, of which 87,394 (~243 h) have been reviewed by an expert. Of these 87,394 10-second clips, 10,293 (~29 h) contain at least one focal or non-focal vocalization with high-resolution onset and offset markers. Associated label files are saved as HDF5 files with the same base filename as the corresponding audio file. The total dataset size, including both audio and associated label files, is 269 GB. The published files use a randomized naming scheme such that the identity of individuals and the recording time of each file cannot be recovered. However, this information can be requested from the CCAS team.

We produced 5-fold stratified 80/20 test/train splits for model training and provided them as tab-separated text files, called manifests, along with the data. Furthermore, we provide a set of train/test manifest files in which recordings of 4 selected individuals are excluded from training and used exclusively for testing. This enables users to assess the generalization performance of trained models to unseen individuals. See Figure 3 for sample count statistics per class.

**Figure 3.**
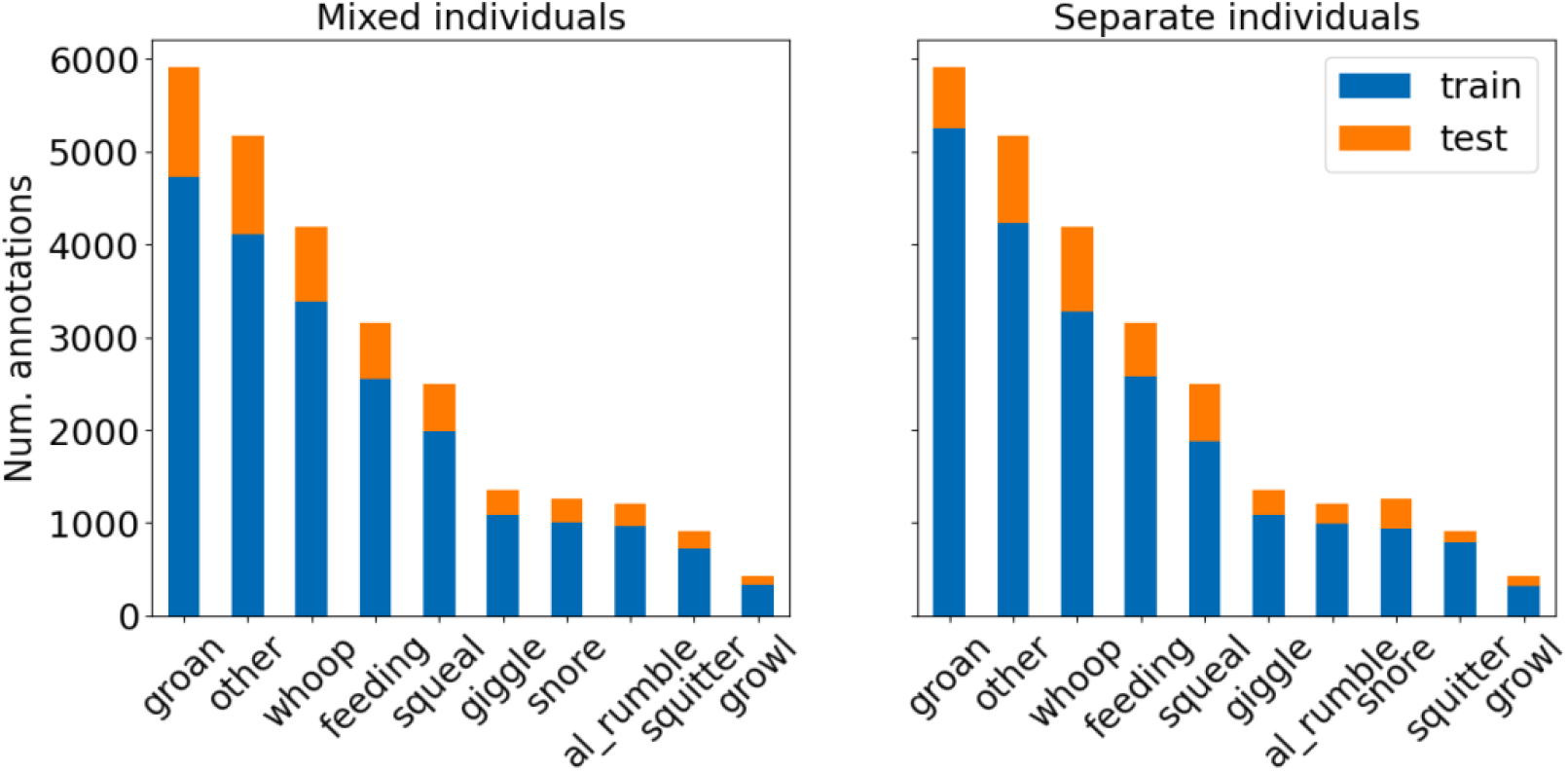
Distribution of the annotations in the train and test splits for both sets of manifest files. The left subplot illustrates the manifest files in which all individuals are represented both in the train and test set. The right subplot corresponds to the manifest files set in which individuals do not overlap in the train and test sets.

We share the dataset as an archive file with three subfolders. The first subfolder contains the wav audio files, the second one the hdf5 label files, and the last one the manifest files. The archive is freely available on the Edmond data repository together with a README file describing the data.^33^

Each 10s clip that has been audited (n = 87,394) has an associated label file with an identical name, which is empty if no vocalizations were found. Each non-empty label file contains the fields described in Table 2. The temporal distribution of all provided classes is shown in Figure 4.

**Table 2.** Description of the label file fields.

| h5 fields | Example | Description |
| --- | --- | --- |
| start_frame_lbl | 15400 | Starting sample point of the vocalization |
| end_frame_lbl | 20000 | Ending sample point of the vocalization |
| foc | 0 | Whether the vocalization is focal (1) or not (0) |
| lbl | “squeal” | Vocalization type |
| lbl_cat | 3 | Integer representing the vocalization type |
| start_time_lbl | 4.30 | Start time of the vocalization in seconds |
| end_time_lbl | 5.20 | End time of the vocalization in seconds |

**Figure 4.**
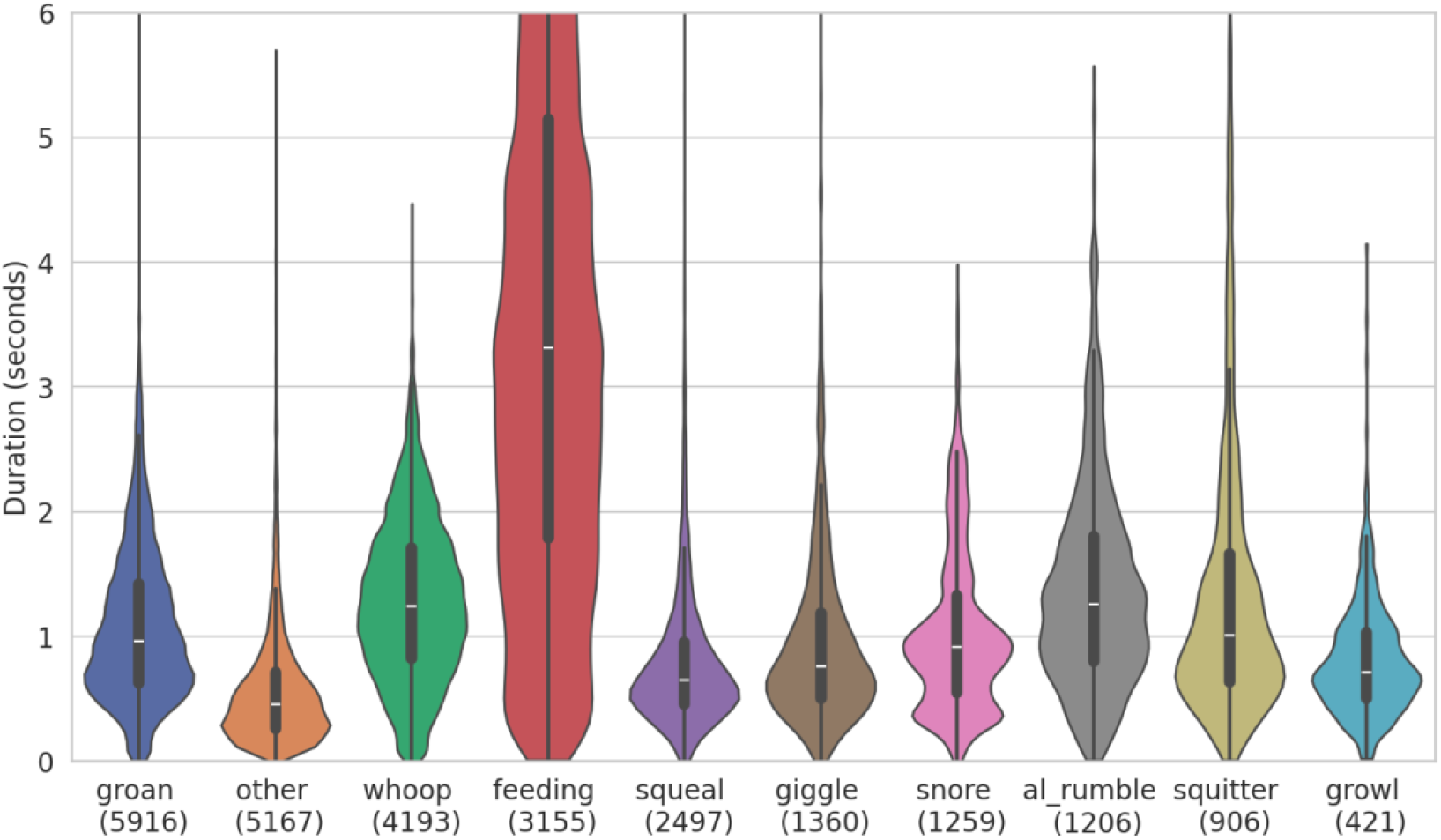
Duration distributions of all sound event classes. These distributions are highly skewed and not Gaussian, so the mode and median are both lower than the mean. The number below the class name represents the number of annotated events within that class.

## Technical Validation

### Benchmark animal2vec model

We provide a benchmark for sound event detection and classification using *animal2vec*.^34^ *animal2vec* is a self-supervised two-step training scheme^35^ using a transformer model^36^ as a basis, designed for sparse bioacoustic data. In the first step, called pretraining, the model learns from all available data without the need for ground truth labels. In self-supervised learning, this is done by artificially creating a supervision signal used to solve a regression or classification task. In *animal2vec*, the scheme includes a ‘student’ and a ‘teacher’ network that share their weights. The student receives a temporally masked version of the full input that the teacher receives. The student’s task is then to regress on the teacher’s output. This way, the model learns to extract information from the data without requiring any labels. The second step, called finetuning, uses available ground truth information to solve a classification task. All reported results here are from a complete pre-trained and fine-tuned model, where we pretrained on all available data and finetuned on the first cross-validation split. *animal2vec* is an open-source tool available at the official *animal2vec* GitHub repository^33^.

### Benchmark animal2vec performance

To evaluate model performance, we use precision, recall, and average precision (AP). Precision assesses the quality of the model by quantifying the accuracy of its positive predictions. In contrast, recall measures completeness, determining the model’s ability to successfully identify all actual true instances within the data. These two metrics inherently exist in a trade-off, meaning that an improvement in one usually results in the degradation of the other.

We visualize this relationship using precision-recall (PR) curves (Figure 5). These curves plot precision against recall across a range of model likelihood thresholds, which represent the model’s varying confidence levels. To compress a precision-recall curve into a single, comprehensive value, we calculate the AP. The AP serves as a robust estimator of the area under the PR curve.

**Figure 5.**
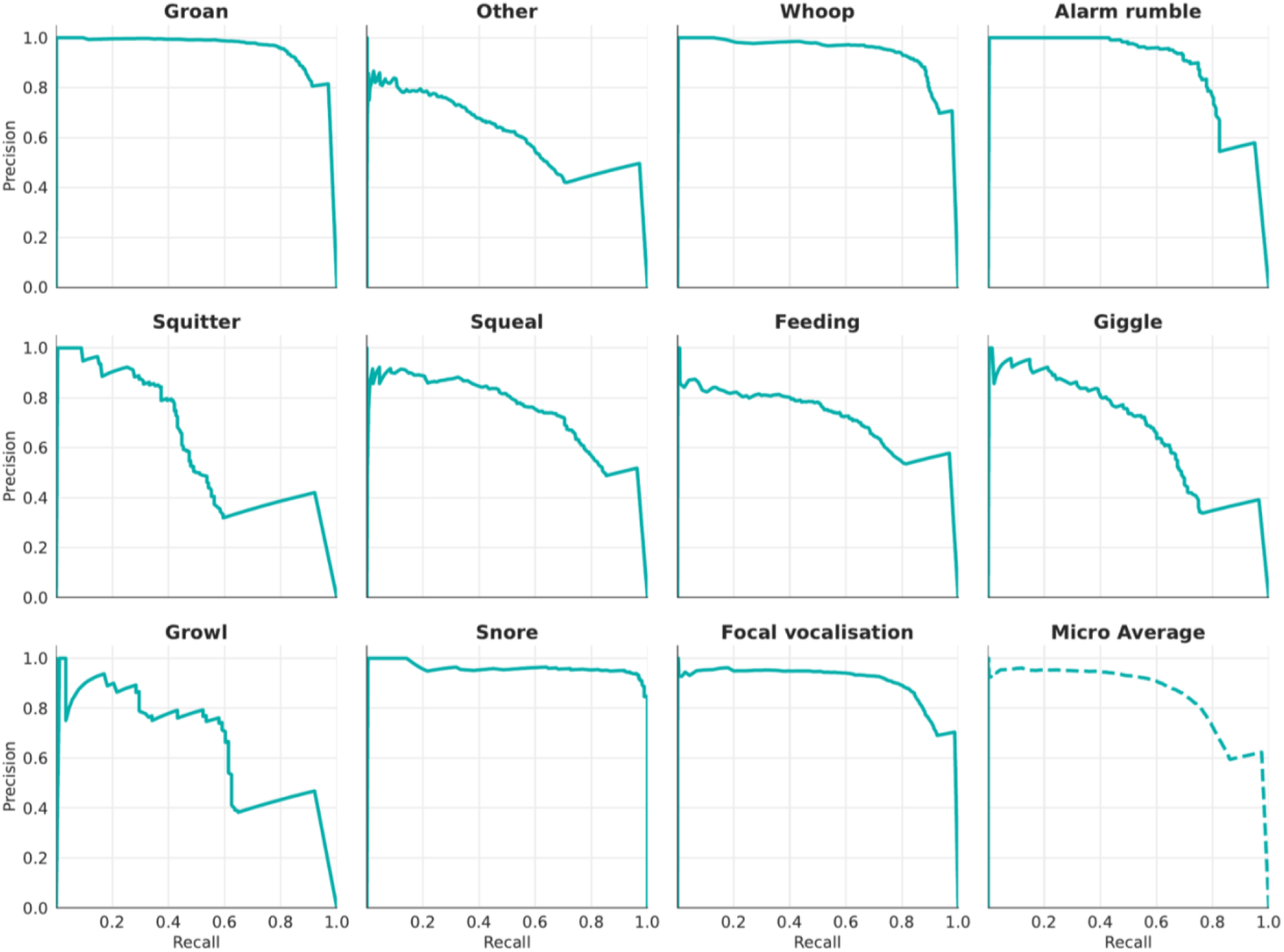
Precision-Recall curves evaluating *animal2vec* classification performance for each class in HyenaSET.

Furthermore, the derived AP scores are aggregated on a per-instance basis to provide meaningful insights. While the *animal2vec* framework natively generates predictions on a granular, frame-by-frame level, we use the post-processing step proposed by the model’s authors.^34^ To identify event boundaries, a sliding average-pooling window is applied to the model’s likelihood output, which is then binarized using a fixed threshold. Final scalar likelihoods are assigned to predicted spans with an Intersection-over-Union (IoU) greater than 0.5. This procedure transforms the per-frame predictions into instance-level metrics (one prediction for one event). The AP scores for each class are shown in Table 3.

**Table 3.**
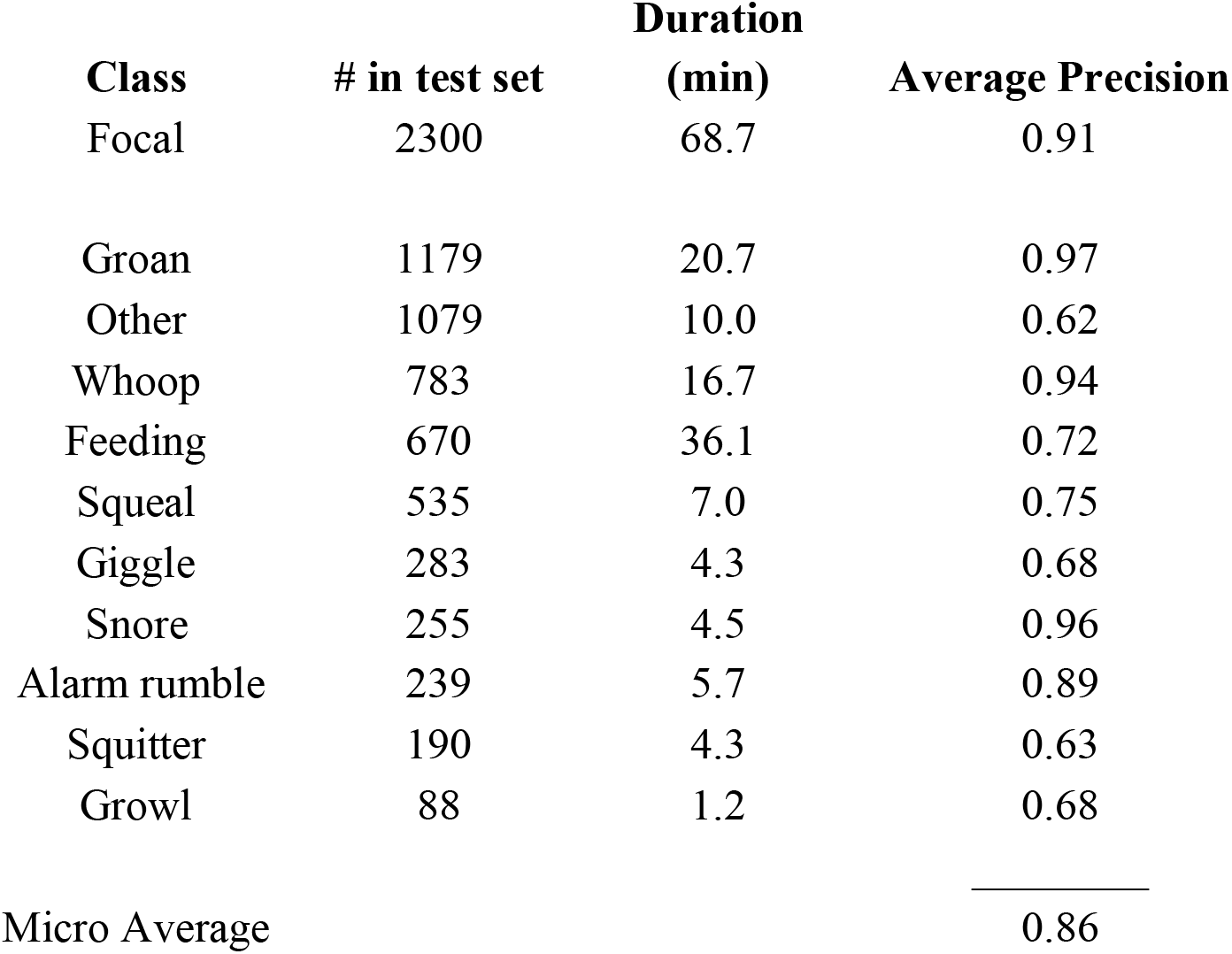
Average Precision scores, number of samples in the test split, and duration in minutes for all classes.

## Data Availability

We share the dataset as an archive file on the Edmond data repository together with a README file describing the data at https://doi.org/10.17617/3.8ZSP3J. We encourage anyone interested in biological analysis of these data to contact the authors.

## Code Availability

The code used to produce the manifest files, train the animal2vec model, and produce the results table is available in the *animal2vec* repository on Github (https://github.com/livingingroups/animal2vec). The exact commands used to implement animal2vec for analysis of these data are shared in the description of the data in the Edmond repository.

## Acknowledgements

We thank the Kenya Wildlife Service, the Kenyan Wildlife Research and Training Institute, the Narok County Government, the Kenyan National Committee on Science, Technology, and Innovation, the Mara Conservancy, and Brian Heath for permissions to collect this data. We would like to thank Dee White, David Nchoko, Kayla Fowler, and the rest of the Mara Hyena Project field team for their help deploying tracking collars on the hyenas. We are also grateful to Marius Faiss, Chi Hsin Chen, Melike Somtürk, Ema Nesic, Ilkim Koc, and Valentin Daschner for their data annotation efforts.

## Author Contributions

Jana M. Woerner - methodology, data curation, writing (original draft), writing (review & editing)

Céline Angonin - data curation, visualization, writing (original draft), writing (review & editing) Andrew S. Gersick - conceptualization, funding acquisition, methodology, writing (review & editing)

Kay E. Holekamp - methodology, resources, writing (review & editing) Mark P. Johnson - methodology, software, writing (review & editing) Marsden Onsare - methodology, writing (review & editing)

Malit O. Pioon - methodology, writing (review & editing)

Julian Schäfer-Zimmermann - data curation, formal analysis, software, visualization, writing (original draft), writing (review & editing)

Frants H. Jensen - conceptualization, funding acquisition, methodology, software, supervision, writing (review & editing)

Ariana Strandburg-Peshkin - conceptualization, funding acquisition, methodology, supervision, writing (original draft), writing (review & editing)

Eli D. Strauss - methodology, data curation, supervision, writing (original draft), writing (review & editing)

## Competing Interests

We declare no competing interests.

## Funding

This work was supported by Human Frontier Science Program award RGP0051/2019 to AS-P and KEH and National Science Foundation grants OISE1853934 and IOS1755089 to KEH. AS-P received additional funding from the Gips-Schüle Stiftung, the Zukunftskolleg at the University of Konstanz, the Max Planck Society, and the Deutsche Forschungsgemeinschaft (DFG, German Research Foundation) under Germany’s Excellence Strategy – EXC 2117 – 422037984.

## Notes

https://edmond.mpg.de/previewurl.xhtml?token=70833c7d-b7a5-44ad-9f8d-b4993a19bc36

### Competing Interest Statement

The authors have declared no competing interest.

### Summary of Updates

This version of the manuscript has been revised to update author affiliations

